# Three-dimensional contact point determination and contour reconstruction during active whisking behavior of the awake rat

**DOI:** 10.1101/2020.03.09.983387

**Authors:** Lucie A. Huet, Hannah M. Emnett, Mitra J. Z. Hartmann

**Affiliations:** Department of Mechanical Engineering, Northwestern University, Evanston, Illinois, 60208; Department of Biomedical Engineering, Northwestern University, Evanston, Illinois, 60208

**Author notes:** **Corresponding Author:** Mitra J. Z. Hartmann, Mechanical Engineering Department, Northwestern University, 2145 Sheridan Road, Evanston, IL 60208.

**Keywords:** Keywords, Vibrissa, whisker, trigeminal, tactile, haptic, active sensing, touch

## Abstract

The rodent vibrissal (whisker) system has been studied for decades as a model of active touch sensing. There are no sensors along the length of a whisker; all sensing occurs at the whisker base. Therefore, a large open question in many neuroscience studies is how an animal could estimate the three-dimensional location at which a whisker makes contact with an object. In the present work we simulated the exact shape of a real rat whisker to demonstrate the existence of a unique mapping from triplets of mechanical signals at the whisker base to the three-dimensional whisker-object contact point. We then used high speed video to record whisker deflections as an awake rat whisked against a peg and used the mechanics resulting from those deflections to extract the contact points along the peg surface. A video shows the contour of the peg gradually emerging during active whisking behavior.

## 1. Introduction

Rats and mice can obtain detailed tactile information by rhythmically sweeping their whiskers back and forth against surfaces and objects in the environment, a behavior called “whisking.” They can use this whisker-based tactile information to determine an object’s location, size, orientation, and texture [1–6].

How rats achieve these tasks is still an open question, especially given that a whisker is simply a cantilever beam with no sensors along its length. Numerous neurophysiological and behavioral studies have specifically investigated how a rodent might use a single whisker to determine the location of a vertical peg [5, 7–20]. Studies have shown that although barrel cortex is required for peg localization [13], knowledge of instantaneous whisker position is not [14]. To date, however, studies have not been able to determine the exact physical cues that the animal might use for peg localization, in part because they have been limited to a two-dimensional (2D) analysis of whisker motion and object contact, with the third dimension sometimes attributed to whisker identity [8, 9].

Complementing the biological literature, several studies in the field of robotics have investigated the problem of whisker-object contact point determination in three dimensions [21–23, 24]. These studies have focused on the use of quasistatic mechanical signals – three reaction forces and three moments at the whisker base – to infer the three-dimensional (3D) whisker-object contact point. Early work showed that using all six signals at the base of a stiff antenna was sufficient to determine the 3D contact point location [22, 24]. and more recent work has shown that in many cases only three of the six mechanical signals are actually required [23]. Which three of the six mechanical signals are needed depends on the intrinsic shape of the whisker, that is, whether it is cylindrical or tapered, straight or curved [23].

The present work brings together these two fields of research to ask how the mechanics associated with whisker bending could allow an actively whisking rat to determine the 3D location of whisker object contact. We used high speed video to record 3D deflections of the “gamma” whisker as an awake rat whisked against a peg, and then simulated the mechanical signals generated by a whisker of that exact shape. We first show that there exists a unique mapping from a triplet of mechanical signals at the whisker base to the 3D location of the whisker-object contact point. We then show that this mapping can be used to extract the 3D contact points, and thus the contour, of a peg placed in front of the animal. These results will generalize across all biologically-realistic whisker shapes with a few exceptions, addressed in the discussion.

## 2. Results

### 2.1. Problem statement: mapping mechanical signals at the whisker base to the 3D whisker-object contact point location

When a rat whisks against an object, as depicted in Figure 1A, the contact point between the whisker and the object is denoted by the coordinates (r_wobj_, θ_wobj_, φ_wobj_) relative to the whisker basepoint, where the subscript “wobj” stands for whisker-object [25]. The whisker’s deflection causes reaction forces and moments (torques) at the whisker base, denoted as *F_x_, F_Y_, F_Z_, M_x_, M_Y_*, and *M_Z_*. The force *F_x_* is called the “axial” force because it acts directly along the whisker’s long axis at the whisker base. The axial force is positive when it pulls the whisker directly out of the follicle and negative when it pushes the whisker directly into the follicle. The forces *F_y_* and *F_z_* are called “transverse” forces, because they act perpendicular (“transversely”) to the whisker at the whisker base. *M_x_* is called the “twisting” moment because it twists the whisker about its long axis, while *M_y_* and *M_z_* are called the “bending” moments because they cause the whisker to bend.

**Figure 1.**
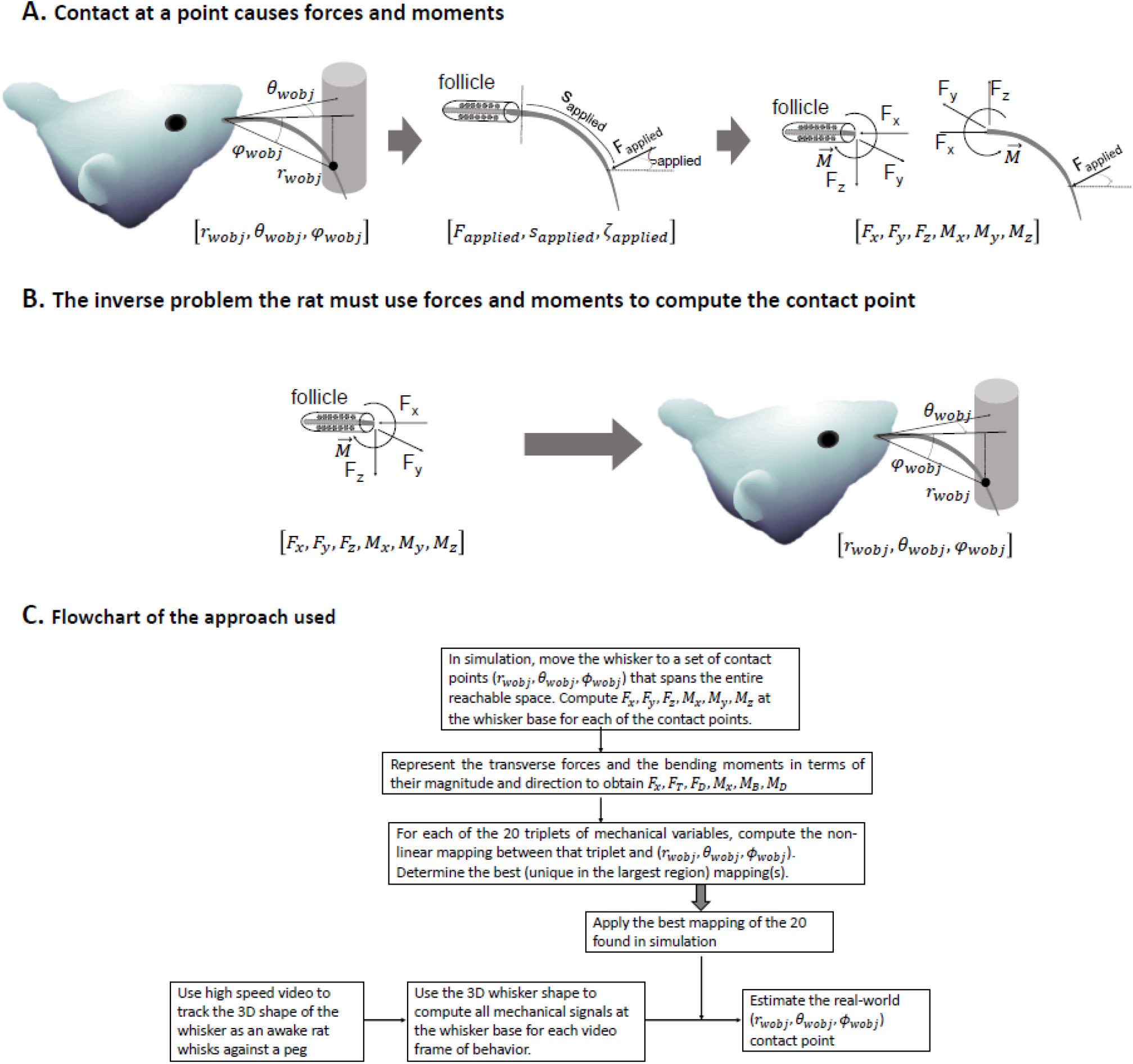
When a rat whisks against an object, forces and moments are generated at the whisker base; the rat must use these forces and moments to determine the 3D contact point location. **(A)** The 3D contact point between the whisker and the object is denoted as r_wobj_, θ_wobj_, and φ_wobj_. This contact point exerts a force on the whisker (F_applied_), which generates reaction forces and moments at the whisker base. **(B)** The inverse problem: the rat’s nervous system must perform the inverse of the process depicted in (A). It must use the forces and moments at the whisker base as inputs to deduce the contact point location. **(C)** A flowchart that describes the approach taken to determine the non-linear mappings that could allow the rat to solve this inverse problem. The 20 mappings indicated in the center box are listed in Supplementary Figure S1.

During exploratory behavior, the rat must solve the inverse problem, illustrated in Figure 1B: it must use the mechanical signals at the whisker base to determine the 3D contact point location. Under the assumption of quasistatic contact, it can be theoretically shown that the six mechanical signals, *F_x_, F_Y_, F_Z_, M_x_, M_Y_*, and *M_Z_* are always sufficient to uniquely determine the 3D contact point location (r_wobj_, θ_wobj_, φ_wobj_) [22, 24].

This theoretical result, however, leaves several important questions unanswered. First, are all six mechanical signals really needed, or might only a subset of them be sufficient to determine the 3D contact point location? Second, assuming that mappings were found that successfully mapped between mechanical signals and contact point, what is the nature of these mappings, and could they be used during real-world exploratory behavior to determine the contours of an object? The answers to these questions directly constrain the neural computations that might permit object localization.

The present work was undertaken to answer these questions, and the procedure used is depicted in the flowchart of Figure 1C. The 3D shape of a real whisker is obtained, and the whisker is then simulated to be deflected to a gridded sampling of contact points across its entire reachable space. For each deflection, the forces and moments at the base of the whisker are computed, and, for convenience, the transverse forces and the bending moments are rewritten in terms of their magnitude and direction. Next, each possible triplet of the six mechanical variables is investigated to determine in which regions it yields a unique mapping to the (r_wobj_, θ_wobj_, φ_wobj_) contact point. The best mapping (i.e., the one that is unique in the largest region) is selected.

In parallel, high speed video is used in behavioral experiments to obtain the 3D shape of the same whisker as an awake rat whisks against a peg. For each video frame the whisker’s deflected shape is used to compute the forces and moments at the whisker base. The best mapping – obtained from the simulation steps described above – is then applied in each video frame to obtain an estimate of the 3D contact point for that frame.

### 2.2 Simulating forces and moments at the base of whisker deflected to all possible positions

Following the procedure depicted in Figure 1C, we began by characterizing the 3D shape of the first whisker in the C-row, traditionally called the “gamma” whisker (Figure 2A). We then simulated deflecting the whisker to a gridded sampling of the 3D point locations it could reach and computed the resulting forces and moments at the whisker base. The set of reachable contact points is depicted as a gray point cloud about the whisker in Figure 2B. For each gray point in Figure 2B, we computed the six forces and moments at the whisker base (*F_x_, F_Y_, F_Z_, M_x_, M_Y_, M_Z_*). We then decomposed the transverse forces and the bending moments into their magnitude and direction:

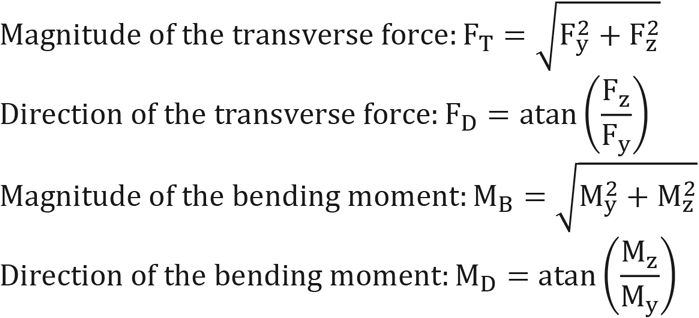

**Figure 2.**
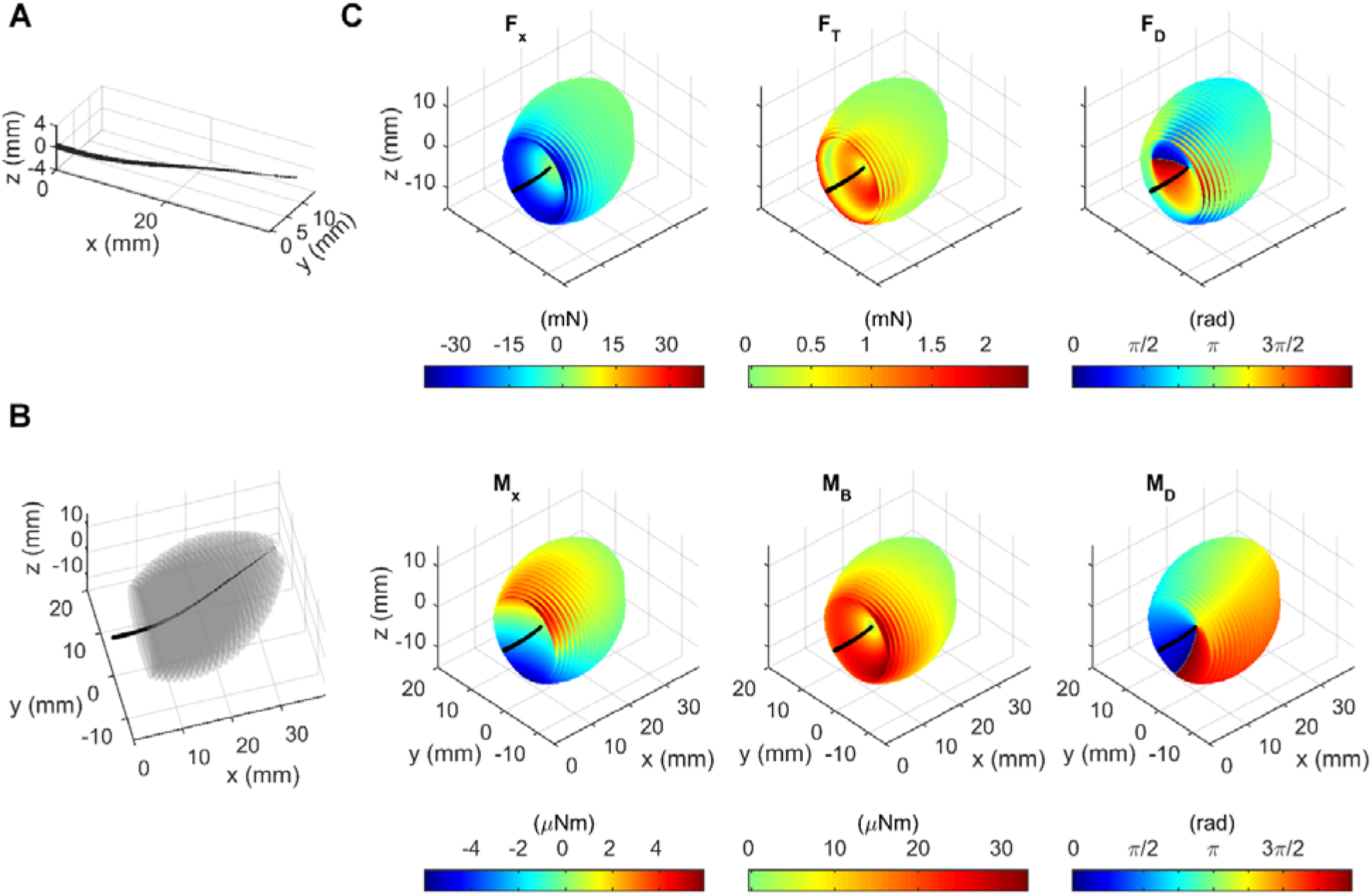
Forces and moments at the whisker base for all possible contact points that the whisker can reach. **(A)** The black line represents the shape of the whisker used in whisker-centered coordinates. The taper in the whisker is for illustration and not to scale. **(B)** The region of contact points that the whisker was able to reach are plotted as a gray cloud around the whisker. The contact points are plotted as a series of translucent gray surfaces; each surface represents a set of reachable contact points at a single radial distance. The translucent surfaces are so close together that they merge into a cloud of points in the shape of an irregular ellipsoid. **(C)** The same whisker and point cloud are shown in each plot at a slightly different angle. The contact points are now colored according to the mechanical variable indicated in each panel. As in (B), the contact points are drawn as surfaces at each radial distance.

These six signals, *F_X_, F_T_, F_D_, M_X_, M_B_*, and *M_D_*, are plotted in the six panels of Figure 2C. Each panel of Figure 2C shows the same black whisker and the same cloud of points as in Figure 2B, but viewed from a different angle for visual clarity. The contact points are colored according to the magnitude of the mechanical signal depicted in the panel. These colored ellipsoid-shaped plots reveal several important trends in the forces and moments at the whisker base.

Two notable trends are that *F_X_, F_T_*, and *M_B_* all exhibit the largest magnitude for proximal contacts and large angles of deflection and that *F_D_* and *M_D_* are always offset 90° from each other. These effects are unsurprising.

The axial force, *F_X_*, has a small region of positive values; in these regions the axial force is pulling the whisker out of the follicle instead of pushing it in. These values occur for distal contacts that are concave forward with small deflections, i.e., the region in which the contact “straightens out” the whisker.

The transverse force, *F_T_*, generally follows the same trends as *M_B_* and *F_X_*, but it also has a hollow “tube” of zero magnitude surrounding the whisker; the “ring” of its bottom end is visible in same relative region where *F_X_* has a large magnitude negative value. This tube occurs when the whisker is bent such that the portion of the whisker local to the contact point is parallel to the y-z plane; in this case the force points entirely in the negative x-direction, resulting in zero *F_T_*. Contact points that deflect the whisker beyond this tube are defined as “large deflection” contacts. This flip into large deflections can also be seen by the sudden 180° change in *F_D_*.

The twisting moment, *M_X_*, exhibits very different trends from the other forces and moments. The magnitude primarily varies in the z-direction rather than radially. This effect occurs because the whisker exhibits more “twist” and therefore greater *M_X_* magnitude as it is deflected out of the x-y plane.

Overall, these trends in forces and moments are similar, but not identical, to those for an idealized, planar, tapered whisker with a parabolic shape [23]. We therefore anticipated that we would see similar results for uniqueness of the mapping between subsets of the variables *F_X_, F_T_, F_D_, M_X_, M_B_, M_D_* and *r_wobj_, θ_wobj_*, φ_*wobj*_.

### 2.3. Unique mappings from triplets of mechanical variables to the 3D whisker-object contact point

Continuing to follow the procedure depicted in the flowchart of Figure 1C, we selected all possible triplets of mechanical signals and tested whether each triplet was sufficient to uniquely determine the 3D location of the whisker-object contact point, *r_wobj_, θ_wobj_*, φ_*wobj*_. Three triplets of mechanical variables were found to meet the criteria for uniqueness: (*F_X_, M_B_, M_D_*), (*M_X_, M_B_, M_D_*), and (*F_X_, F_D_, M_D_*). Criteria for uniqueness can be found in Supporting Information, along with a list of the regions of non-uniqueness for the remaining 19 mappings (Supplementary Table S1).

Each of these successful mappings can be visualized with a set of three colored, solid shapes. However, these visualizations are unintuitive and challenging to understand. To provide intuition for how to visualize a mapping we show an example using the (*F_X_, M_B_, M_D_*) triplet. Figure 3 depicts the gradual construction of the three solids that represent the mapping between (*F_X_, M_B_, M_D_*) and (*r_wobj_, θ_wobj_*, φ_*wobj*_).

**Figure 3:**
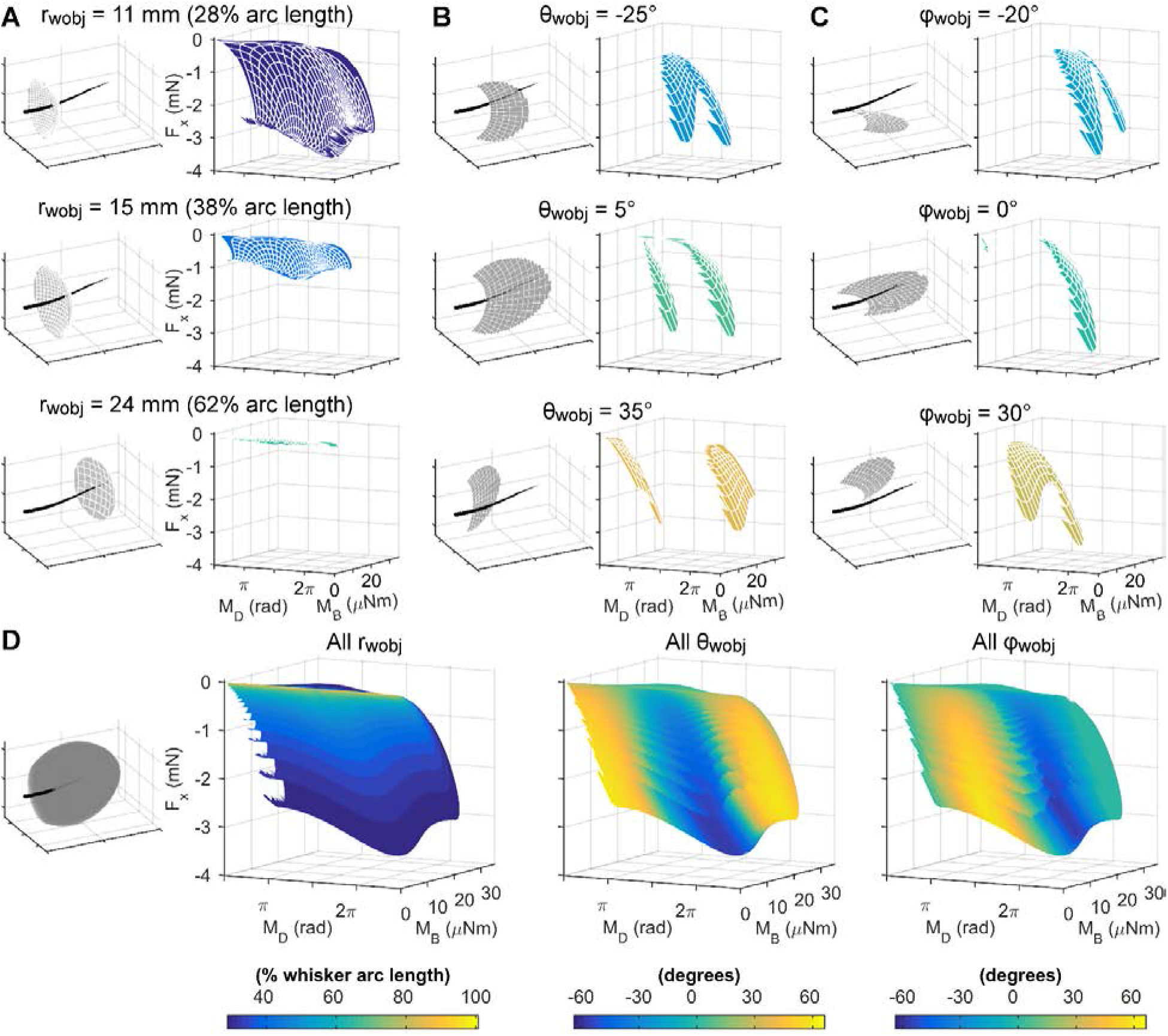
The mappings between mechanical variables and 3D whisker-object contact point can be represented as solids, each represented as series of monochromatic surfaces. **(A)** *Left column:* The whisker is depicted in black, with all reachable contact points at a given radial distance shown as a gray surface. The three rows show contact points for radial distances of 11, 15, and 24 mm. *Right column:* The corresponding F_X_, M_B_, and M_D_ signals for the contact points at each of the radial distances are plotted as monochromatic surfaces. **(B)** *Left column:* The whisker is shown in black, with all the reachable contact points at a given azimuthal angle shown as a gray surface. The three rows show contact points for θ_wobj_ = 25°, 5° and 35 °. *Right column:* The corresponding F_X_, M_B_, and M_D_ signals for the contact points at each of the values of θ_wobj_ are plotted as monochromatic surfaces. **(C)** *Left column:* The whisker is shown in black, with all the reachable contact points at a given elevation angle shown as a gray surface. The three rows show contact points for φ_wobj_ = −20°, 0° and 30 °. *Right column:* The corresponding F_X_, M_B_, and M_D_ signals for the contact points at each of the values of φ_wobj_ are plotted as monochromatic surfaces. **(D)** The monochromatic surfaces for each of the r_wobj_, θ_wobj_, and φ_wobj_ mappings are layered together to form three “solids.” *First column:* All the contact points in gray about the whisker in black. This plot contains the identical points as Fig. 2B. *Second column:* All the monochromatic surfaces for the different r_wobj_ values plotted together in F_X_, M_B_, M_D_ space. The “feathered edge” is an artifact of the discretization of the r_wobj_ values. *Third column:* All the monochromatic surfaces for the different θ_wobj_ values. *Fourth column:* All the monochromatic surfaces for the different φ_wobj_ values.

The three panels in the left column of Fig. 3A show the whisker in black along with all the contact points it could reach at three different radial distances: 11 mm, 15 mm, and 24 mm. These distances correspond to 28%, 38% and 62% of the whisker arc length, respectively.

The three panels in the right column of Fig. 3A show *F_X_, M_B_*, and *M_D_* computed from the contact points at the three different radial distances. In each of these three right panels, the points representing *F_X_, M_B_*, and *M_D_* are connected to form a single continuous surface. Each surface is monochromatic, indicating that all points within that surface are generated from contact points at the same radial distance. The surface corresponding to r_wobj_ = 15 mm is a different color from the surface corresponding to r_wobj_ = 11 mm, and its shape is “shrunk down” on the *F_X_* and *M_B_* axes. Similarly, the surface for *F_X_, M_B_*, and *M_D_* corresponding to r_wobj_ = 24 mm is yet a third color and its shape has shrunk even more in *F_X_* and *M_B_*.

Fig. 3B shows a similar set of plots as Fig. 3A, except that each surface represents a single θ_wobj_ angle, instead of a single value for r_wobj_. The three panels in the left column show gray contact points at three different values of θ_wobj_: −25°, 5°, and 35°, and the three panels in the right column show the corresponding monochromatic surfaces in the *F_X_, M_B_, M_D_* space. Notice that the color scale for θ_wobj_ (Figure 3B) is independent of the color scale for r_wobj_ (Figure 3A).

Fig. 3C shows similar plots for three different values of φ_wobj_: −20°, 0°, and 30°, with corresponding monochromatic surfaces in the *F_X_, M_B_, M_D_* space. Again, notice that the color scale for φ_wobj_ is independent of the color scale for θ_wobj_ and r_wobj_.

Finally, Fig. 3D combines all the surfaces for each geometric coordinate into its own plot. In the left column of Fig. 3D all the contact points plotted as a gray cloud about the whisker; this figure matches Figure 2B. The second column in Fig. 3D shows all the monochromatic surfaces in *F_X_, M_B_, M_D_* space for r_wobj_. Notice that all the different surfaces are nested one inside the other. This set of surfaces represents the mapping “solid,” forming a lookup table to determine r_wobj_. If given three values, *F_X_, M_B_*, and *M_D_* at the base of the whisker for an unknown contact, the color of the solid at that *F_X_, M_B_, M_D_* location determines the radial distance. The solid lookup tables for *θ_wobj_*, and φ_*wobj*_ are shown in the third and fourth columns of Fig. 3D; again these solids are formed by plotting a series of monochromatic surfaces.

As expected, the solids in columns 2 through 4 of Figure 3D all have the same shape; they differ only in coloring. The “feathered edges” most noticeable on the visualization for r_wobj_ are a discretization artifact and do not have any significance. Using a larger number of values for r_wobj_ would cause the feathered edges to coalesce into the identical shape as those for *θ_wobj_*, and φ_*wobj*_. Close visual examination of the solids in Figure 3D reveals an important feature: none of the monochromatic surfaces within any of the solids overlap. The absence of overlap indicates that a single reading of *F_X_, M_B_*, and *M_D_* results in unique values for r_wobj_, *θ_wobj_*, and φ_*woby*_, meaning that the mapping is unique. If any of the monochromatic surfaces overlapped or intersected, then the readings of *F_X_, M_B_*, and *M_D_* at that point of intersection would result in multiple combinations of r_wobj_, *θ_wobj_*, and φ_*woby*_, rendering the mapping non-unique.

Importantly, however, due to human fallibility, visual inspection is necessary but not sufficient to determine if a particular mapping is unique. In addition to visual inspection, we tested uniqueness using neural networks as non-linear function solvers (details provided in Supporting Information). If a neural network could solve for a mapping, then the mapping was unique. Specifically, given (F_X_, M_B_, M_D_) as inputs and (r_wobj_, *θ_wobj_*, φ_*wobj*_) as outputs, the network had to be able to solve for the non-linear function that maps between inputs and outputs with sufficiently small errors. In other words, the neural network effectively generates a “look-up table” for the mapping, and the look-up table must be unique.

We found that – as expected – the look up table generated using (F_X_, M_B_, M_D_) as inputs contained especially high errors in r_wobj_ when the contact point was near the whisker and deflections were very small. These high errors are expected because tiny variations in the forces generated during small angle deflections can cause large changes in r_wobj_ for the estimated contact point. The look-up table did not have sufficient resolution to resolve contact points for these small angle deflections. In future work this issue could be addressed in part by changing the mesh distribution of the contact points, but in the present work we simply excluded plotting contact points that fell within a thin “cone” surrounding the whisker. Specifically, Figures 4 and 5 omit any point that lay less than s_closest_*tan(2°) from the whisker, where s_closest_ is the arc length of the point on the whisker closest to the contact point.

**Figure 4:**
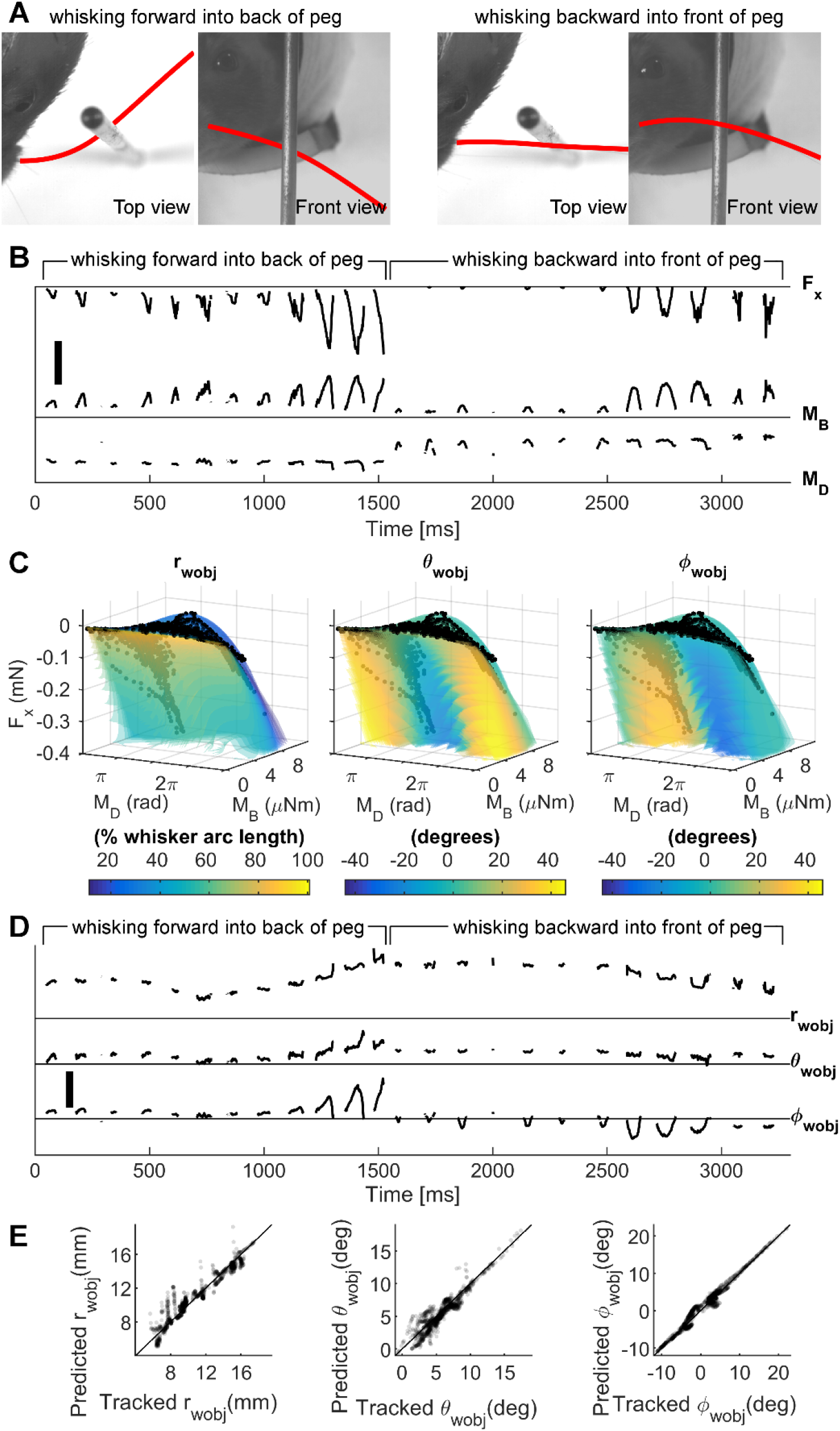
The 3D contact point (r_wobj_, θ_wobj_, φ_wobj_) can be determined from F_X_, M_B_, M_D_ during active whisking. **(A)** Top and front views from the raw video data of a rat whisking forward into the back of the peg (left two images) and backwards into the front of the peg (right two images). The tracked whisker is traced in red. **(B)** The F_X_, M_B_, M_D_ data output by Elastica3D as the rat whisked into the peg. Note the sudden change in M_D_ when the rat switches from whisking forward to whisking backward into the peg. Scale bar: 0.25 mN for F_X_, 10 μNm for M_B_, and 2π radians for M_D_. **(C)** F_X_, M_B_, and M_D_ from B plotted onto the mapping visualizations for r_wobj_, θ_wobj_, and φ_wobj_. These visualizations show the relevant parts of the mappings from Fig. 3D, and the monochromatic layers are plotted as translucent for better visualization. The black dots represent the data from B plotted onto F_X_, M_B_, M_D_ space. The color of the “solid” at the black dots in each of the three plots gives the contact point location coordinates: r_wobj_, *θ_wobj_*, φ_*wobj*_. **(D)** The r_wobj_, θ_wobj_, and φ_wobj_ values given by the mappings in C. Scale bar: 10 mm for r_wobj_, 20° for θ_wobj_ and φ_wobj_. **(E)** Tracked values of r_wobj_, θ_wobj_, and φ_wobj_ plotted against the values of r_wobj_, θ_wobj_, and φ_wobj_ as predicted from the mechanical signals. The dots are translucent to improve visualization. The black diagonal line represents where the dots would lie if the 3D contact points computed from the mechanical signals were perfect.

**Figure 5:**
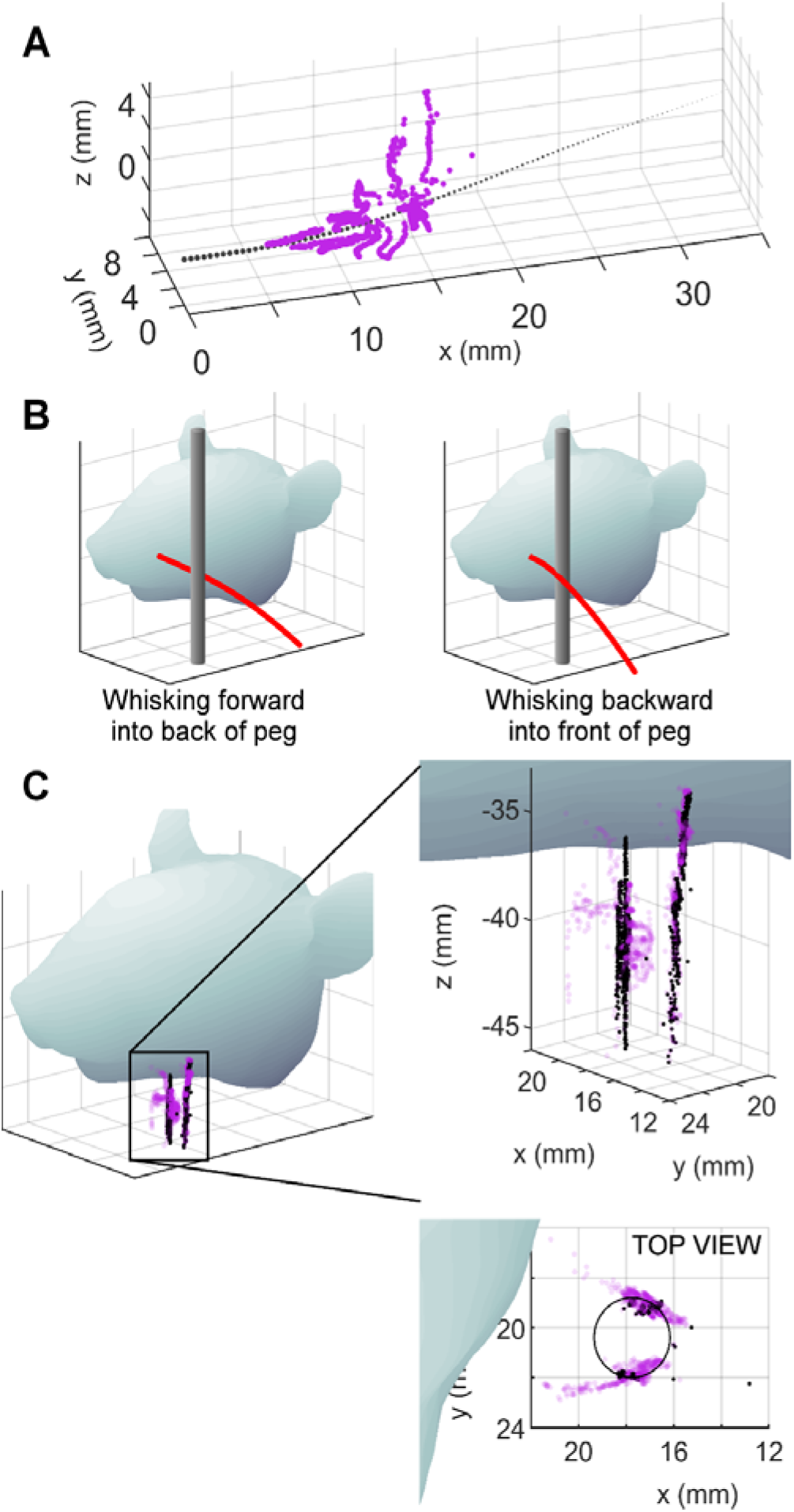
The r_wobj_, θ_wobj_, φ_wobj_ coordinates computed from the mechanical signals at the whisker base can reconstruct the shape of the peg. **(A)** The r_wobj_, θ_wobj_, φ_wobj_ coordinates computed from the mechanical signals are plotted in whisker-centered coordinates. The whisker is the black tapered, dotted line. Each magenta dot represents a 3D contact point on the peg with respect to the whisker. Each dot is computed based on mechanical data obtained from a single video frame (1 ms resolution).These are the identical points as were plotted in Figure 4D, but now plotted in their correct 3D spatial locations relative to the whisker. **(B)** Diagrams of the rat whisking into the peg show where reconstructed contact points are expected. The rat first whisks forward against the back of the peg, and then whisks backwards against the front of the peg. We therefore expect to see reconstructions of both the front and the back of the peg. **(C)** The same view of the rat head as in (B), now including contact points. Tracked (“ground truth”) contact points are shown as black dots. Contact points computed from mechanical signals are shown as transparent magenta dots. Two expanded views of the reconstructed peg are shown to the right. In the side view, the points create two vertical lines: one indicating contact with the back of the peg, and the other indicating contact with the front of the peg. The top view reveals that the largest errors are found in the estimate of radial distance, rather than in estimates of the angles of contact.

### 2.4 Contact point determination in the awake, behaving rat

The mappings described in the previous sections were obtained purely from simulation. The simulations assumed that the whisker was rigidly clamped at its base and underwent ideal, frictionless point-deflections.

It is not at all evident that the mappings obtained from simulation will apply during active whisking behavior of an awake rat. The mapping results shown in Figure 3D could be degraded by many nonlinear effects, including tissue compliance [11] and non-ideal multi-point or sliding contact [26]. The mappings are therefore only of theoretical interest unless the associated lookup tables can be successfully applied to real-world whisking behavior.

To address this important concern we recorded ~3.5 seconds of high-speed video as an awake rat whisked against a vertical peg (2.7 mm diameter). During this particular trial of whisking, the rat first whisked forward against the back of the peg. The whisker then slipped past the peg, and the rat whisked backwards against the front of the peg. Fig. 4A shows the top and front views of the rat whisking forward and backward against the peg with the tracked whisker traced in red.

For each frame of video, we used the tracked 3D shape of the whisker and the tracked location of the 3D whisker-object contact point to compute the mechanical signals *F_X_, M_B_*, and *M_D_* at the whisker base. These mechanical signals, shown in Figure 4B, represent the information we assume the rat has during whisking behavior [25, 27, 28]. Details for finding the 3D whisker shape and contact point location are provided in Supplementary Figure 2.

The three mechanical signals, *F_X_, M_B_*, and *M_B_*, were then used as the inputs to the mapping established in Figure 3D. During this particular trial of whisking, the signals spanned a more limited range than that shown in Figure 3D. Therefore, Figure 4C illustrates only the region of the mappings relevant to this particular whisking trial. Each of the mapping solids in Figure 4C is slightly translucent, so as to show a set of black dots representing the trajectory of the whisker through the F_X_, M_B_, and M_D_ space. This time-varying trajectory is best observed in Supplementary Video 1.

In Figure 4C, the color of each of the solid objects at the location of the black points yields the reading for the estimated contact point location. Using the appropriate look-up table for this mapping, we can obtain predicted values for the contact point (r_wobj_, *θ_wobj_*, φ_*wobj*_) which are plotted as functions of time in Fig. 4D.

The quality of the predicted values for (r_wobj_, *θ_wobj_*, φ_*wobj*_) was evaluated by plotting them against the ground-truth tracked values for (r_wobj_, *θ_wobj_*, φ_*wobj*_); results are shown in Figure 4E. Data points for each measurement at 1 msec intervals are represented by a black dot. If they fall on the black diagonal line, the predicted value exactly matches the tracked value. The fact that the points generally lie close to the diagonal line shows the high quality of the mapping between mechanical signals and the 3D contact point.

As expected, predictions from the look-up table show the greatest errors for the variable r_wobj_. These errors result from limitations in the resolution of the mappings for small deflections: very small changes in force can cause large jumps in predictions for r_wobj_.

### 2.5 Reconstructing the shape of a peg from the 3D contact points

To further evaluate the quality of the mapping between (*F_X_, M_B_, M_D_*) and (r_wobj_, *θ_wobj_*, φ_*wobj*_), we performed a coordinate transformation to reconstruct the shape of the peg in the laboratory frame. Figure 6A shows the whisker in black with the predicted 3D contact points plotted in whisker-centered coordinates around the whisker. Whisker-centered coordinates are somewhat unintuitive, so to understand this figure, imagine deflecting the whisker to each of the contact points in turn. These deflections are identical to those that occurred as a result of the rat’s active whisking against the peg. The points are dense near the proximal section of the whisker because the rat happened to whisk against the peg using the proximal portion of its whisker.

The next step was to convert these contact points to the laboratory frame, schematized in Figure 5B. Figure 5C is the identical schematic, but the peg has now been replaced with whisker-object contact points. The black dots in Figure 5C are the ground-truth contact points tracked directly from video, while the magenta dots represent the reconstruction of the peg as determined from the mappings. Notice that the “ground truth” as described here still contains tracking error; it is not the outline of the peg.

As expected, the black dots form two vertical lines: one on the back of the peg and one on the front of the peg. The reconstructed contact points are best visualized in Supplementary Video 2. The magenta dots match relatively well with the tracked peg points; however, the top view (inset) reveals that the contact point estimates are somewhat mis-matched with respect to r_wobj_. This result is consistent with the results shown in the first column in Fig. 4E, depicting the goodness of fit for r_wobj_. Supplementary Figure S3 gives an explanation for how errors in the peg reconstruction in both Figures 4 and 5 result from limitations in mapping resolution.

## 3. Discussion

### 3.1. Approaches towards finding the 3D whisker-object contact point

Unlike an insect antenna, a mammalian whisker has no sensors along its length: all mechanical sensing occurs at the whisker base. Therefore, a long-standing question is how an animal could determine the 3D location at which a whisker contacts an object. Previous proposed solutions have fundamentally relied on two general approaches.

The first approach involves measuring rates of change of one or more mechanical variables at the whisker base, most typically the bending moment. Specifically, it can be shown that the rate of change of bending moment is related to the radial distance of contact [5, 11, 29–33]. The second approach, used in the present work, involves combining multiple geometric variables [11] or mechanical signals [5, 33] in a nonlinear manner, independent of the rates of change of these signals. The major advantage of the second approach over the first is that it is history independent [34]. In other words, the contact point can be calculated at each instant of time, and the calculation does not depend on the trajectory of the whisker on the surface of the object.

A recent behavioral study strongly suggested that animals make at least some use of the second approach [5]. In these experiments, mice whisked either against a rigid peg at a radial distance far from the whisker base or against a compliant peg at a radial distance close to the whisker base. The two peg positions were precisely chosen so as to ensure that the rate of change of bending moment was the same for both. Nonetheless, the mice could still easily distinguish between the peg locations. These results indicate that rodents do not use a localization strategy that relies exclusively on measuring the time rates of change of the bending moment.

Which mechanical signals can be combined so as to uniquely determine the 3D whisker-object contact point? The answer depends on the shape of the whisker, including its taper and intrinsic curvature. Given the standard approximation that a whisker is planar, linearly tapered, and is either straight or has a parabolic intrinsic curvature, the signals F_X_, M_B_, and M_D_ can be shown to uniquely map to r_wobj_, θ_wobj_, and φ_wobj_ [23]. Whiskers in the real world, however, do not have this idealized geometry, and the goal of the present study was therefore to determine which triplet(s) of mechanical signals could generate a unique mapping for a real whisker, with non-planar, non-parabolic curvature.

### 3.2 Mappings for a real, non-idealized whisker

The “gamma” whisker used in the present study curved out of the plane and did not have a parabolic curvature. For this particular whisker, three triplets of mechanical variables were found to generate unique mappings to the 3D contact point: (*F_X_, M_B_, M_D_*), (*M_X_, M_B_, M_D_*), and (*F_X_, F_D_, M_D_*). We chose to focus on the first triplet because it is associated with the most general mapping: simulations have shown that it provides a unique mapping to (r_wobj_, θ_wobj_, φ_wobj_) for all tapered whiskers, regardless of whether they are straight or have intrinsic parabolic curvature [23]. We first computed the signals F_X_, M_B_, M_D_ at the base of the gamma whisker during a bout of whisking behavior and then showed that these signals can be used to successfully estimate 3D contact point location (Figure 5 and Supplementary Video 2). In other words, the results show that the unique mappings found in simulation for an idealized whisker can extend to a real-world, non-planar whisker with non-parabolic curvature.

Generalizing, these results demonstrate that a rodent could use mechanical information entering the follicle from a single whisker to determine the 3D contact point location in whisker-centered coordinates. The mapping between mechanical signals and 3D geometry exists independent of whisking phase or velocity, that is, the mapping remains constant for a given whisker across all whisks. In addition, the 3D contact point can be determined at all points in times during a deflection, which means that the approach can continuously solve for 3D contact location as the whisker sweeps across an object surface or even deflects against a compliant object. We note that this work is a direct and natural extension of the two-dimensional mapping from (F_X_, M_B_) to (r_wobj_, θ_wobj_) described in previous work [33].

### 3.3 Model limitations and their effects on the mapping

The two main limitations to the whisker model used, Elastica3D, are that it assumes quasistatic and frictionless conditions.

The quasistatic assumption will affect the accuracy of the mappings only if the dynamic response of the whisker significantly interferes with the mechanical response due to bending. This interference can easily be avoided if the whisker maintains contact with an object long enough for dynamic effects to damp. Recent work has shown that rats maintain contact with an object for 20-50 msec [35, 36], which is exactly long enough for dynamic effects to dissipate [37].

Friction could potentially have a larger effect on mapping accuracy. In the absence of friction, each 3D contact point is associated with one and only one deflected whisker shape and one set of forces and moments at the whisker base. When friction is included, however, a single contact point location could result in multiple deflected whisker shapes and the forces and moments will depend on the history of contact. A larger coefficient of friction will have a greater effect on the mappings and could significantly reduce mapping uniqueness [5, 38, 39]. Frictional effects will be an important topic for future investigation.

### 3.4. Expected generalization across whiskers and whisker shapes

The present study has focused on a single gamma whisker, with a single set of parameters. What parameter changes affect the mappings and their uniqueness? The mappings will undoubtedly change for whiskers that have different shapes, but the question is how large these changes will be, and whether the F_X_, M_B_, M_D_ mapping will retain its uniqueness.

It is reasonable to assume that almost all naturally occurring whiskers will be tapered [40–43], be largely (but not entirely) planar [44, 45], and have a curvature that is mostly well-described by either a quadratic or cubic equation [44–46]. Trimming the tip of the whisker, as might occur through natural damage or barbering, would have no effect on mapping uniqueness. The only change will be that the whisker cannot reach as large a region of space. Similarly, changing Young’s modulus will scale all mechanical variables at the whisker base but not affect mapping uniqueness. Changing the radius of the whisker base, or changing the whisker arc length, will have no effect on mapping uniqueness, provided that the ratio of the base radius to tip radius remains constant and r_wobj_ is measured as a fraction of the whisker arc length (instead of in terms of absolute distance).

Changing the radius slope of the whisker (defined as the difference between the base and tip radii, divided by the arc length) will change the base-to-tip radius ratio, which will affect the mappings in highly nonlinear and often unpredictable ways. However, the mapping will remain unique for the F_X_, M_B_, M_D_ triplet as long the whisker retains sufficient taper. If the taper is very slight, the mapping will theoretically be unique, but the resolution required to distinguish (r_wobj_, θ_wobj_, φ_wobj_) might be so high as to be impractical [23]. If the whisker is cylindrical, the mapping will be non-unique [23].

Based on these considerations, we expect unique mappings to hold for nearly all naturalistic whisker shapes. Mappings will be non-unique if there is very large in-plane curvature (e.g., if the whisker curves so much that it doubles back on itself), if there is very large out-of-plane curvature (e.g., if the whisker is a corkscrew), or if the whisker has no taper. With the exception of these cases, the problem of mapping “uniqueness” may be more accurately posed as a problem of mapping resolution. For example, Figure 4E indicates that high error occurs when r_wobj_, θ_wobj_ are small. Future work will help resolve the origin of these errors and shed light on the extent to which the F_X_, M_B_, M_D_ mapping will generalize across arbitrary whisker shapes.

### 3.5. Implications for object localization and robotics

During object localization, deformations of the whisker cause a wealth of mechanoreceptors to deform within the follicle [47–51] in response to whisker bending. The whisker deformations are most easily represented in whisker-centered coordinates. The mechanoreceptors transduce these deformations to electrical signals, which are transmitted to primary sensory neurons in the trigeminal ganglion (Vg). For an animal to then localize an object in head-centered coordinates, more central levels of the trigeminal pathway would need to integrate information about the whisker’s position and orientation on the face [12, 17, 18, 25]. Alternatively, a recent study suggests that the mapping between mechanical signals in whisker-centered coordinates and the location of an object in head-centered coordinates may also be unique [52]. This implies that the animal would not need information about the whisker orientation but instead could localize using only whisker-centered information, reducing central level processing.

The results of the present work pave the way to develop robots that can perform accurate 3D contour extraction [34]. Provided that the signals obtained from the base of artificial whiskers contain sufficient information about the relevant mechanics, they could be combined in a highly-nonlinear mapping to determine the 3D contact points along the contours of an object in whisker-centered coordinates. Later, these contact points can then be converted to robot-centered coordinates. Because hardware models inherently include effects such as collisions, vibrations, and friction, they may be particularly useful to help neuroscientists constrain processing at more central levels of the trigeminal system. Finally, these models could help inform physics simulations of active whisking in real-world conditions.

## 4. Methods

Ethics statement: All procedures involving animals were approved in advance by the Institutional Animal Care and Use Committee (IACUC) of Northwestern University.

### 4.1 Quantifying vibrissa motion and shape

All whiskers on the left side of the face of a female Long-Evans rat (3-6 months) were trimmed except for the Gamma whisker. The rat was body-restrained, and two orthogonally-mounted high speed video cameras recorded whisking behavior against a vertical peg.

The cameras were Photron 1024PCI monochrome cameras (1,000 fps, shutter speed 1/3000 second, lenses Nikon AF Micro-Nikkor 60 mm) and were mounted equal distances (~60 cm) from the rat. One camera obtained a top-down (“bird’s eye”) view and the other obtained a front-on view. Each pixel (58 μm) was matched between the two cameras using a 2×2 mm^2^ checkerboard grid. In the top-down camera view, the whisker was tracked using open source software “whisk” [53]. The whisker was manually tracked in the front-on camera view. Because the two views shared the same pixel scaling and were positioned perfectly orthogonal to each other, merging was straightforward. Pixels along the x-axis were matched between the two views, and the tracked whiskers and contact points (x-y data in the top view, x-z data in the front-on view) were combined into 3D representations [54].

In each frame, the whisker basepoint position, the whisker’s angles of emergence [45, 46], and the 3D contact point location were tracked. The basepoint position and the whisker’s angles of emergence were filtered at 85 Hz to smooth tracking jitter. The cutoff 85 Hz was chosen because it both eliminated tracking noise while preserving the peaks of the whisker motion. Filtering at higher frequencies generally resulted in noisier contact point estimates; this was a gradual effect as the filtering frequency was increased. Filtering at significantly lower frequencies did not accurately capture the whisker’s bending as assessed by matching the raw tracked data with the filtered estimates.

### 4.3 Generating Forces from Behavioral Data and for Mappings

The whisker shape obtained in the section “Quantifying vibrissa motion and shape” above was run through two different simulations: one to obtain forces and moments at the whisker base during active whisking, and one to find forces and moments for the entire reachable space of the whisker. The same parameters were used in both simulations. The radius at the base was 100 μm, similar to that of a C2 whisker [55], and a base radius to tip radius ratio of 15 was used [41, 43], giving the tip a radius of 6.67 μm. We assumed a typical value for Young’s modulus of 3 GPa [55] and for Poisson’s ratio, 0.38 [56].

From the 3 seconds of data of a rat actively whisking into a peg, we used the tracked undeflected whisker shape, 3D contact point at all frames of contact, and position and orientation of the whisker at all frames of time. For each frame of contact, we used the whisker’s position and orientation to place the contact point in whisker-centered coordinates. We then used this contact point in whisker-centered coordinates as input to Elastica3D (more information available in Supplementary Methods and at https://github.com/SeNSE-lab/DigitalRat) with the undeflected tracked whisker to find the forces and moments at the base of the rat whisker.

In order to generate force and moment data for the mappings, we deflected the whisker to points distributed over nearly the entire space of contact points the whisker could reach. Spherical coordinates were used to describe the contact point locations. The radial distance (r_wobj_) ranged from 6 mm (30% of the whisker arc length) to 20 mm (100% of the whisker arc length) in millimeter increments. The azimuthal angle (θ_wobj_) ranged from −65° to +65° in single degree increments, and the elevation angle (φ_wobi_) ranged from −60° to +60° in single degree increments. The simulation deflected the whisker to all the resulting contact point locations. If Elastica3D could not converge to a solution for a contact point, meaning the whisker would slip off the contact point, then the point was discarded. The forces and moments for all the remaining contact points were then recorded for use in the mappings.

### 4.4 Mapping uniqueness

The uniqueness of each of the 20 mappings using different triplet combinations of forces and moments at the whisker base was determined using the same methods described in [23]. In order to be unique, a mapping had to be found unique by both visual inspection methods and by a neural network. More details can be found in Supporting Information.

## Supporting Information Legends

**Supplementary Video 1. The 3D contact point (r_wobj_, θ_wobj_, ϕ_wobj_) can be determined from the signals F_X_, M_B_, M_D_ during active whisking. (Top)** The signals F_X_, M_B_, and M_D_ vary with time as the rat whisks against the peg. The sudden change in MD occurs when the rat switches from whisking forward to whisking backward against the peg. Scale bar: 0.25 mN for F_X_, 10 μN-m for M_B_, and 2π radians for M_D_. **(Middle)** Each of the three panels shows the mapping between the signals F_X_, M_B_, and M_D_ (variables on the axes) and one of the three geometric coordinates r_wobj_, θ_wobj_, and ϕ_wobj_ (represented with a colormap). Each mapping is a solid and is drawn as a set of semi-transparent monochromatic layers to aid visualization. In each panel, the trajectory of the black dot represents the changing values of the F_X_, M_B_, and M_D_ signals shown in the *Top* subplot of the video. At each instant of time, the color at the location of the black dot in the three panels indicates the 3D contact point coordinates. **(Bottom)** The geometric coordinates (r_wobj_, θ_wobj_, ϕ_wobj_) generated by the mappings shown in the *Middle* subplot. Scale bar: 10 mm for r_wobj_, 20° for θ_wobj_ and ϕ_wobj_.

**Supplementary Video 2. The r_wobj_, θ_wobj_, ϕ_wobj_ coordinates computed from the mechanical signals at the whisker base can be used to reconstruct the shape of the peg. (Top left)** This panel shows a still image of the rat’s head, the peg location, and the tracked whisker (red). This still image can be compared with the videos in the other panels, all of which show different views of the peg “reconstruction” over time. **(Top center)** The time-varying 3D whisker-object contact point (r_wobj_, θ_wobj_, ϕ_wobj_) computed from the mechanical signals is plotted in whisker-centered coordinates. The whisker is the black, tapered, dotted line. Each magenta dot represents a 3D contact point on the peg with respect to the whisker. Each dot is computed based on mechanical data obtained from a single video frame (1 ms resolution). These are the identical points plotted in all other video panels, but are now plotted in their correct 3D spatial locations relative to the whisker. **(Top right)** Top-down view and front-on view of the rat as it whisks against the peg. The rat first whisks forward against the back of the peg, and then whisks backwards against the front of the peg. The whisker is traced in red for those frames in which it is in contact with the peg. **(Bottom row)** All videos in the bottom row show the gradual reconstruction of the peg’s contour based on computing contact points from the mechanical signals at the whisker base. Because the rat whisks both forward and backwards against the peg, reconstructions are observed for the peg’s front and back sides. Contact points directly tracked from the video (“ground truth”) are shown as black dots. Contact points computed from mechanical signals are shown as transparent magenta dots. **(Bottom left and center)** The left panel shows an isometric view of the rat’s head along with the time-varying whisker-peg contact points, while the center panel shows an expanded view of the contact points. Note that the contact points form two distinct vertical lines, indicating contact with the front and the back of the peg. **(Bottom right)** Reconstruction of the peg contour in top-down and front-on views, matching the videos directly above. The top-down view reveals that the largest errors are found in the estimate of radial distance, rather than in estimates of the angles of contact.

